# Crystal structure of the SARS-CoV-2 non-structural protein 9, Nsp9

**DOI:** 10.1101/2020.03.28.013920

**Authors:** D. R. Littler, B. S. Gully, R. N. Colson, J Rossjohn

## Abstract

Many of the proteins produced by SARS-CoV-2 have related counterparts across the Severe Acute Respiratory Syndrome (SARS-CoV) family. One such protein is non-structural protein 9 (Nsp9), which is thought to mediate both viral replication and virulence. Current understanding suggests that Nsp9 is involved in viral genomic RNA reproduction. Nsp9 is thought to bind RNA via a fold that is unique to this class of betacoronoaviruses although the molecular basis for this remains ill-defined. We sought to better characterise the SARS-CoV-2 Nsp9 protein and subsequently solved its X-ray crystal structure, in an apo-form and, unexpectedly, in a peptide-bound form with a sequence originating from a rhinoviral 3C protease sequence (LEVL). The structure of the SARS-CoV-2 Nsp9 revealed the high level of structural conservation within the Nsp9 family. The exogenous peptide binding site is close to the dimer interface and impacted on the relative juxtaposition of the monomers within the homodimer. Together we have established a protocol for the production of SARS-CoV-2 Nsp9, determined its structure and identified a peptide-binding site that may warrant further study from the perspective of understanding Nsp9 function.

## Introduction

Severe acute respiratory syndrome coronavirus 2 (SARS-CoV-2) is comprised of a large single stranded positive polarity RNA genome that acts as messenger RNA after entering the host. Post infection, the ssRNA encodes two open reading frames, produced through host ribosomal frameshifting that transcribes two polyprotein products. The polyprotein products are subsequently cleaved into 27 viral proteins by internally encoded proteases. Further processing of the polyprotein releases an RNA-polymerase along with several non-structural proteins that facilitate RNA synthesis and may play a role in the enveloping process but are not included in the viral coat.

Treatment of infections caused by betacoronoaviruses have focused on three therapeutic strategies; vaccination with the spike glycoprotein of the SARS-Cov-2 envelope (Wrapp *et al.*, 2020), and small-molecule targeting of conserved viral enzymes (*e.g.* the Mpro protease (Zhenming *et al.* 2020) (Yang *et al.*, 2005) and the RNA-polymerase (Yan *et al.* 2020). Nevertheless, some of the betacoronaviral non-structural proteins appear important for viral replication within SARS-CoV and influence pathogenesis (Frieman *et al.*, 2012). Despite their close homology between viruses, such non-structural proteins remain of interest as they may have conserved roles within the viral lifecycle of SARS-CoV-2.

During infection of human cells, SARS-CoV Non-structural protein 9 (Nsp9_SARS_) was found to be important for replication (Frieman *et al.*, 2012). Homologs of the Nsp9 protein have been identified in numerous coronaviruses including SARS-Cov-2 (Nsp9_COV19_), human coronavirus 229E (Nsp9_HcoV_), avian infectious bronchitis virus (Nsp9_IBV_), porcine epidemic diarrhea virus (Nsp9_PEDV_) and porcine delta virus (Nsp9_PDCoV_). Nsp9_SARS_ has been shown to have modest affinity for long oligonucleotides with binding thought to be dependent on oligomerisation state (Egloff *et al.*, 2004) (Sutton *et al.*, 2004). Nsp9_SARS_ dimerises in solution via a conserved α-helical ‘GxxxG’ motif. Disruption of key residues within this motif reduces both RNA-binding (Sutton *et al.*, 2004) and SARS-CoV viral proliferation (Frieman *et al.*, 2012). The mechanism of RNA binding within the Nsp9 protein family is not understood as these proteins have an unusual structural fold not previously seen in RNA binding proteins (Egloff *et al.*, 2004) (Sutton *et al.*, 2004). The fold’s Greek-key motif exhibits topological similarities with OB-fold proteins and small protein B but such vestiges have proven insufficient to provide clear insight into Nsp9 function (Egloff *et al.*, 2004). As a consequence of the weak affinity of Nsp9_SARS_ for long oligonucleotide stretches it was suggested that the natural RNA substrate may instead be conserved features at the 3’ end of the viral-genome (the stem-loop II RNA-motif) (Ponnusamy *et al.*, 2008). Furthermore, potential direct interactions with the co-factors of the RNA polymerase have been reported (Chen *et al.*, 2017). However, it remains to be determined how the oligonucleotide-binding activity of Nsp9 proteins promote viral replication during infection.

The sequence of Nsp9 homologues are conserved amongst betacoronoaviruses, yet there remains the potential for functional differences in different viruses. Nsp9_COV19_ exhibits 97% sequence identity with Nsp9_SARS_ but only 44% sequence identity with Nsp9_HCoV._ The structure of the HCoV-229E Nsp9 protein suggested a potential oligomeric switch induced upon the formation of an intersubunit disulfide bond. Here, disulfide bond formation shifts the relative orientation of the Nsp9 monomers, which was suggested to promote higher-order oligomerisation (Ponnusamy *et al.*, 2008). The resultant rod-like higher order Nsp9_HCoV_ assemblies had increased affinity for the RNA oligonucleotides. Cysteine mutants of Nsp9_HCoV_ that are unable to produce the disulfide displayed reduced RNA-binding affinity (Ponnusamy *et al.*, 2008). The observation of a redox-induced structural switch of Nsp9_HCoV_ led to the hypothesis that Nsp9_HCoV_ may have a functional role in sensing the redox status of the host cell (Ponnusamy *et al.*, 2008). While the “redox-switch” cysteine responsible for oligomer formation in Nsp9_HCoV_ is conserved amongst different viral Nsp9 homologues the higher-order oligomers were not observed for Nsp9_SARS_ (Ponnusamy *et al.*, 2008). Because of these potential differences between Nsp9 proteins we sought to further characterise the nature of Nsp9_COV19_.

## Results

### Expression and purification of the SARS-CoV-2 Nsp9 protein

The Nsp9 protein from SARS-CoV-2 (Nsp9_COV19_) was cloned and recombinantly expressed in *E. coli*. The expression construct included an N-terminal Hexa-His tag attached via a rhinoviral 3C-protease site. Following Ni-affinity chromatography Nsp9_COV19_ was further purified via size-exclusion chromatography to yield > 95% pure and homogeneous protein. Nsp9_COV19_ eluted from gel filtration columns with the apparent molecular weight of a dimer suggesting that, as with other Nsp9 proteins, Nsp9_COV19_ is an obligate homodimer. The N-terminal tag was removed prior to any biochemical experiments via overnight digestion with precision protease, as reported for Nsp9_SARS_ (Sutton *et al.*, 2004).

### Nucleotide binding of Nsp9COV19 protein

The affinity of viral Nsp9 homologues for oligonucleotides has a range of binding affinities reported, some of which are dependent on oligomerisation state and nucleotide length, ranging from 20-400 μM (Zeng *et al.*, 2018). We therefore sought to assess the potential for Nsp9_COV19_ to bind to fluorescently labelled oligonucleotides using fluorescence anisotropy. Preliminary experiments were performed under conditions similar to those previously identified for Nsp9_SARS_ (Sutton *et al.*, 2004). Oligonucleotide affinity was very limited under our assay conditions (Fig. 1). Indeed, protein concentrations up to 200 μM of Nsp9_COV19_ did not result in saturated binding and thus indicated an incredibly low affinity *K*_D_, or no affinity for these oligonucleotides at all under these assay conditions.

**Fig. 1.**
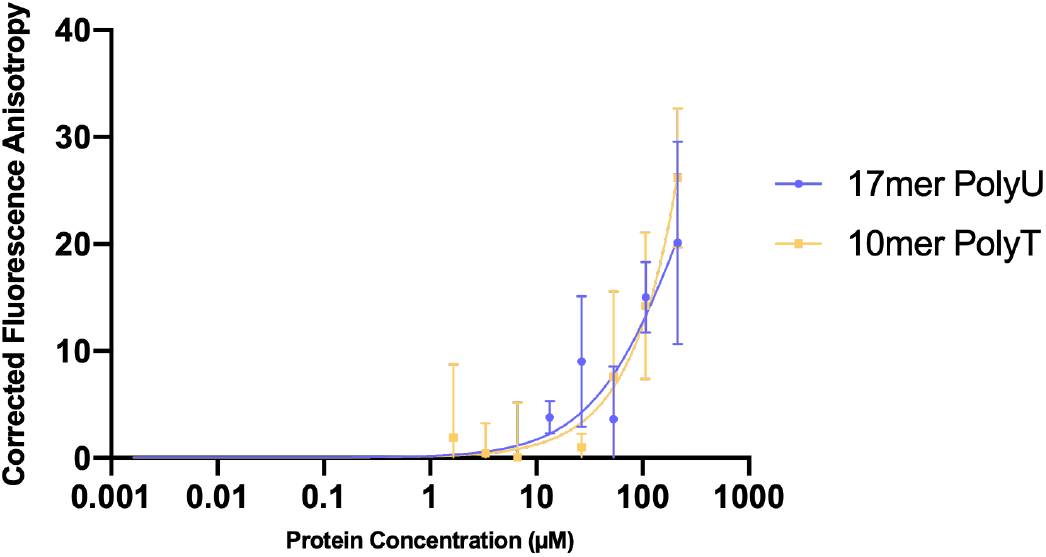
Nsp9_COV19_ nucleotide-binding assay. Fluorescence polarization anisotropy assays were used to examine the possibility that Nsp9_COV19_ could bind to labelled 17-mer and 10-mer single stranded oligonucleotides. The plot shows corrected anisotropy for each Nsp9_COV19_ protein concentration, error bars represent the SD from the mean of triplicate measurements after 60 minutes incubation.

### Crystal structure of apo-Nsp9_COV19_

We next determined the structure of apo-Nsp9_COV19_ (Table 1). The apo-Nsp9_COV19_ structure aligned closely to that of Nsp9_SARS_ (R.M.S.D of 0.57Å over 113 C_α_, Fig. 2A-C) (Egloff *et al.*, 2004). Like other Nsp9 homologues it exhibits an unusual fold that is yet to be observed outside of coronaviruses (Sutton *et al.*, 2004). The core of the fold is a small 6-stranded enclosed β-barrel, from which a series of extended loops project outward (Fig 2A). The elongated loops link the individual β-strands of the barrel, along with a projecting N-terminal β-strand and C-terminal α1-helix; the latter two elements make up the main components of the dimer interface (Fig. 3A). Two loops project from the open-face of the barrel: the β2-3- and β3-4-loops are both positively charged, glycine rich, and are proposed to be involved in RNA-binding. The only protrusion on the enclosed barrel-side is the β6-7-loop; the C-terminal half of the β7-strand is an integral part of the fold’s barrel-core but its other half extended outward to pair with the external β6-strand and create a twisted β-hairpin, cupping the α1-helix and interacting with subsequent C-terminal residues.

**Table 1.**
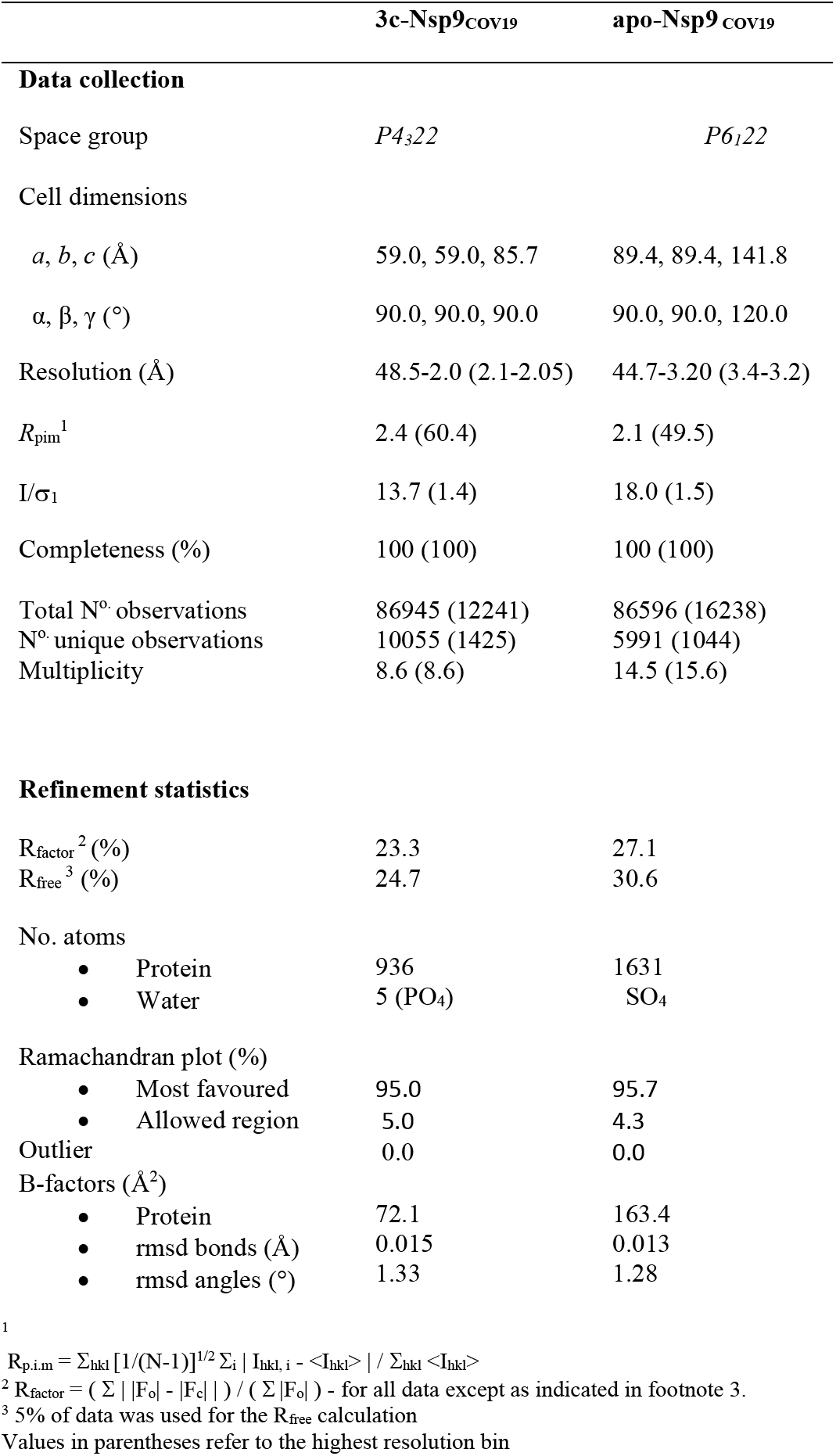
Data collection and refinement statistics

**Fig. 2.**
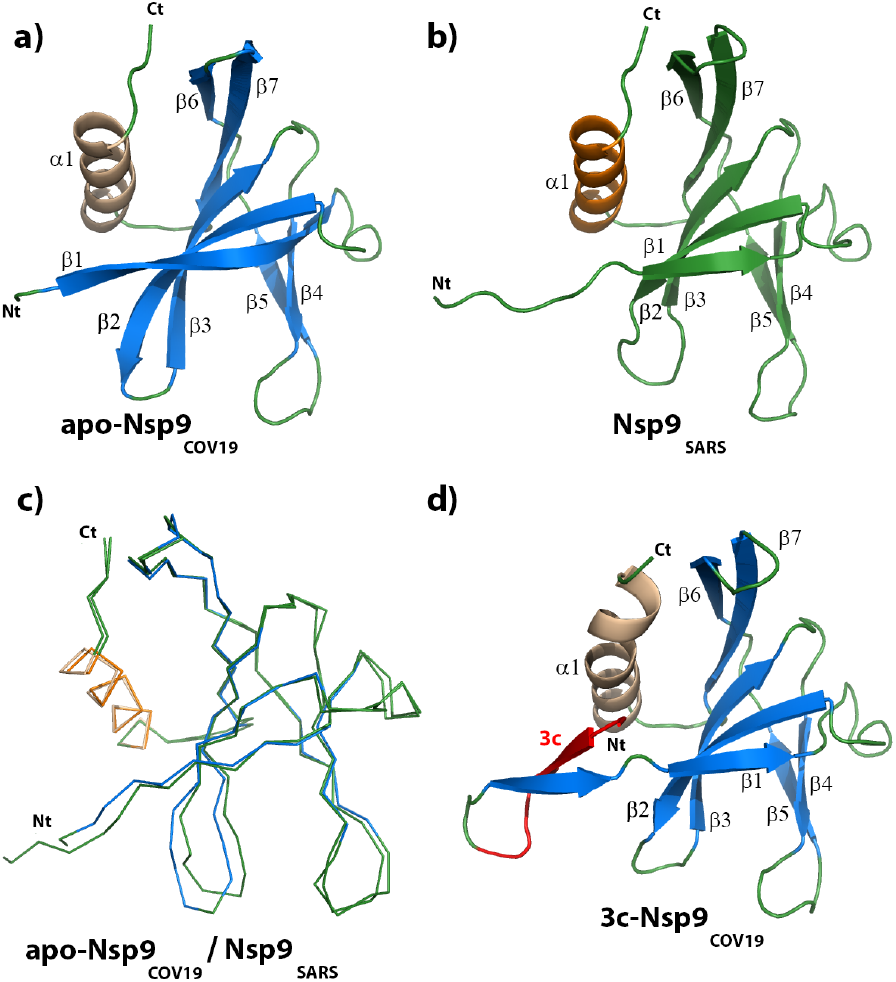
apo-Nsp9_COV19_ is structurally similar to Nsp9_SARS_. Cartoon representation of the monomeric units of **a)** apo-Nsp9_COV19_ **b)** apo-Nsp9_SARS_ (Sutton *et al.*, 2004) and **c)** a backbone alignment of the two structures. The COV19 structures are colored with β-strands in *marine* and the α-helix in *wheat;* the SARS structures are in *teal* and *orange* respectively. **d)** the bound peptide is highlighted in *red*.

**Fig. 3.**
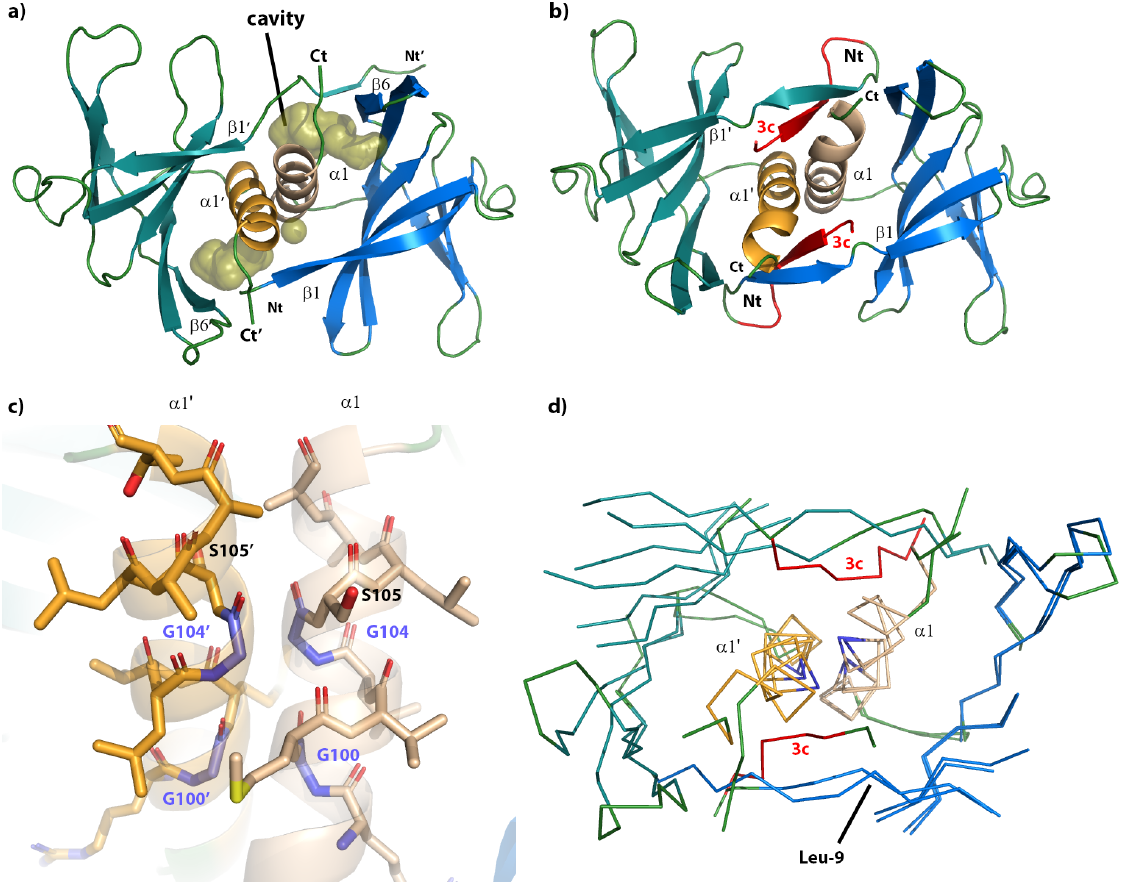
Peptide binding in Nsp9_COV19_ alters the dimer interface. Top-down views of the dimer interface highlighting the interaction helices for **a)** unbound Nsp9_COV19_ in which the surface of the hydrophobic interface cavity is displayed labelled **b)** an equivalent representation of peptide-occupied 3c-Nsp9_COV19_ dimer. **c)** Stick representation of the GxxxG protein-protein interaction helices at the dimer interface for apo-Nsp9_COV19_. **d)** Cα backbone overlay of the Nsp9_COV19_ interface in the apo and peptide-occupied states. The GxxxG motif residues are colored *light purple*.

The arrangement of monomers within Nsp9-dimers is well-conserved in different viruses and is maintained within Nsp9_COV19_ (R.M.S.D of 0.66Å over 226C_α_ compared to the dimeric unit of Nsp9_SARS_). The main component of the intersubunit interaction is the self-association of the conserved GxxxG protein-protein binding motif (Fig. 3C) that allowed backbone van der Waals interactions between interfacing copies of the C-terminal α1-helix (Hu *et al.* 2017). Here Gly-100 of the respective parallel α1-helices, formed complementary backbone van der Waals interactions. These interactions were replicated after a full helical turn by Gly-104 of the respective chains, thereby forming the molecular basis of the Nsp9_COVID19_ dimer interface (Fig. 3C). The 2-fold axis that created the dimer ran at a ~15 ° angle through the GxxxG motif allowing the 14-residue helix to cross its counterpart (Fig. 4A), the N-terminal turns of the helix were relatively isolated, only making contacts with counterpart protomer residues. In contrast, the C-terminal portions were encircled by hydrophobic residues, albeit at a distance that created funnel-like hydrophobic cavities either side of the interfacing helices (Fig. 3A). Strands β1, β6 and the protein’s C-terminus served to provide a ring of residues that encircled the paired helices. The first 10 residues of Nsp9_COV19_ exchanged across the dimer-interface to form a strand-like extension of β1 that ran alongside β6 from the other protomer (Fig. 3A). The interaction these strands made did not appear optimal, indeed the remaining four C-terminal residues projected sideways across the dimer interface, inserting between the two strands while contributing a hydrophobic backing to the main helix.

**Fig. 4.**
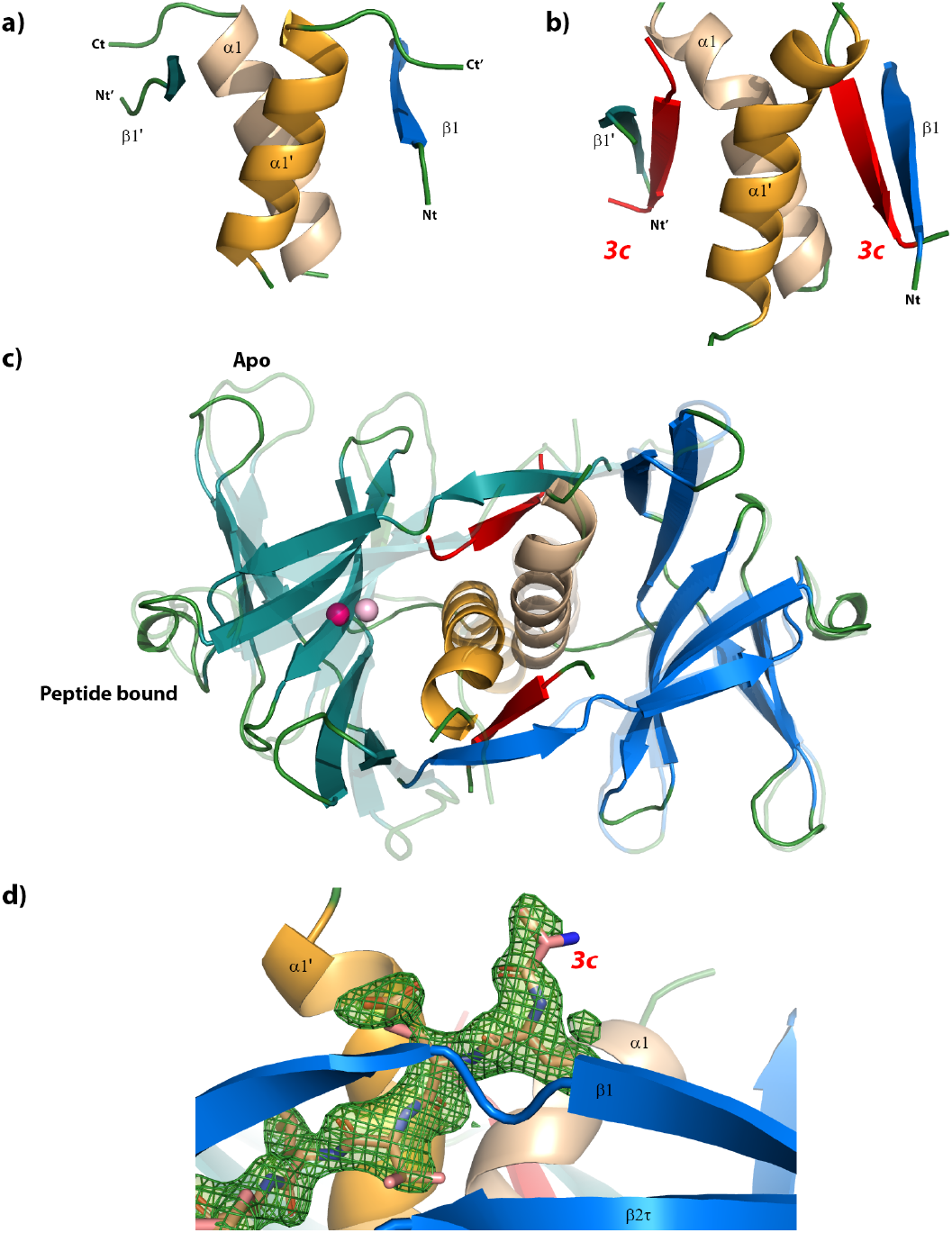
Movements within the GxxxG motif. A side-view of the N- and C-terminal structural elements at the dimer interface are shown for the **a)** apo-and **b)** peptide-occupied forms. **c)** Overlay of the Nsp9_COV19_ dimer in the apo and peptide-bound forms indicating respective shifts in subunit orientation. The center-of-mass of the nonaligned subunit is depicted with a *light-pink* and *dark-pink* point respectively. **d)** Unbiased omit map contoured at 3.2σ near the hydrophobic cavities into which the exogenous bound peptide was refined.

### Extraneous peptides occupy the hydrophobic cavities of Nsp9

In a separate crystallisation experiment we determined the structure of Nsp9_COV19_ that included the N-terminal tag together with a rhinoviral 3C protease sequence (termed 3C-Nsp9_COV19_). The 3C-Nsp9_COV19_ crystal form diffracted to 2.05 Å resolution in space group P4_3_22, and had 1 molecule within the asymmetric unit, with the dimer being created across the crystallographic 2-fold axis.

Unexpectedly the high-resolution structure of 3C-Nsp9_COV19_ diverged from that of the apo-Nsp9_COV19_ (R.M.S.D 0.86 Å for the monomer and 2.23 Å when superimposing a dimer). The 3C sequence folded-around either side of the paired intersubunit helices to fill two funnel-like hydrophobic cavities (Fig. 2D, 3B, 4C, D). Namely, 3C residues LEVL, inserted into the opposing cavities either side of the dimer interface and ran parallel to the paired GxxxG motif. Moreover, the 3C sequence formed additional β-sheet interactions with the N-terminus of the protein from the other protomer (Fig. 3B). To accommodate the 3C residues the N-terminal strand residues moved outward by ~1.6 Å (residues 6-10). This movement allowed the N-terminus to increase the number of β-sheet interactions it formed with β6’. The β-barrel core of the fold remained unchanged but the increase in interactions between β1 and β6’ served to exclude the C-terminus, prompting residues 106-111 to condense into a bent extension of the α-helix (Fig. 4A, B). The subtle structural changes near the interacting GxxxG motifs (Fig. 3D) are amplified at the periphery of the dimer resulting in ~6 Å shift in the β-barrel core (Fig. 4C)

### Conserved cavity residues accommodate a peptide backbone

When comparing apo-Nsp9_COV19_ with 3C-Nsp9_COV19_ the point where the N-terminal interface strand diverges is near Leu-9 (Fig. 3D). Within the apo form it makes van der Waals interactions with the sidechains of Met-101, Asn-33 and Ser-105; this latter serine is important as it immediately follows the conserved protein-binding motif (^100^**G** MVL**G** S^105^), while also specifically interacting with Gly104’ from the opposing protomer. Within the 3C-Nsp9_COV19_ structure the extraneous LEVL residues insert at this point (Fig. 5B), the hydrophobic side chains clasp either side of Ser-105 and allowed its hydroxyl group to form backbone hydrogen bonds to the glutamate within the extraneous sequence (Fig. 5B). Meanwhile the C-terminal Leucine from the extraneous residues inserted behind the α-helix of the other protomer (Fig. 5B). Cumulatively these changes allow for a ~5° rotation of the protomer subunits about the 2-fold axis compared to apo-Nsp9_COV19_ (Fig. 4C).

**Fig. 5.**
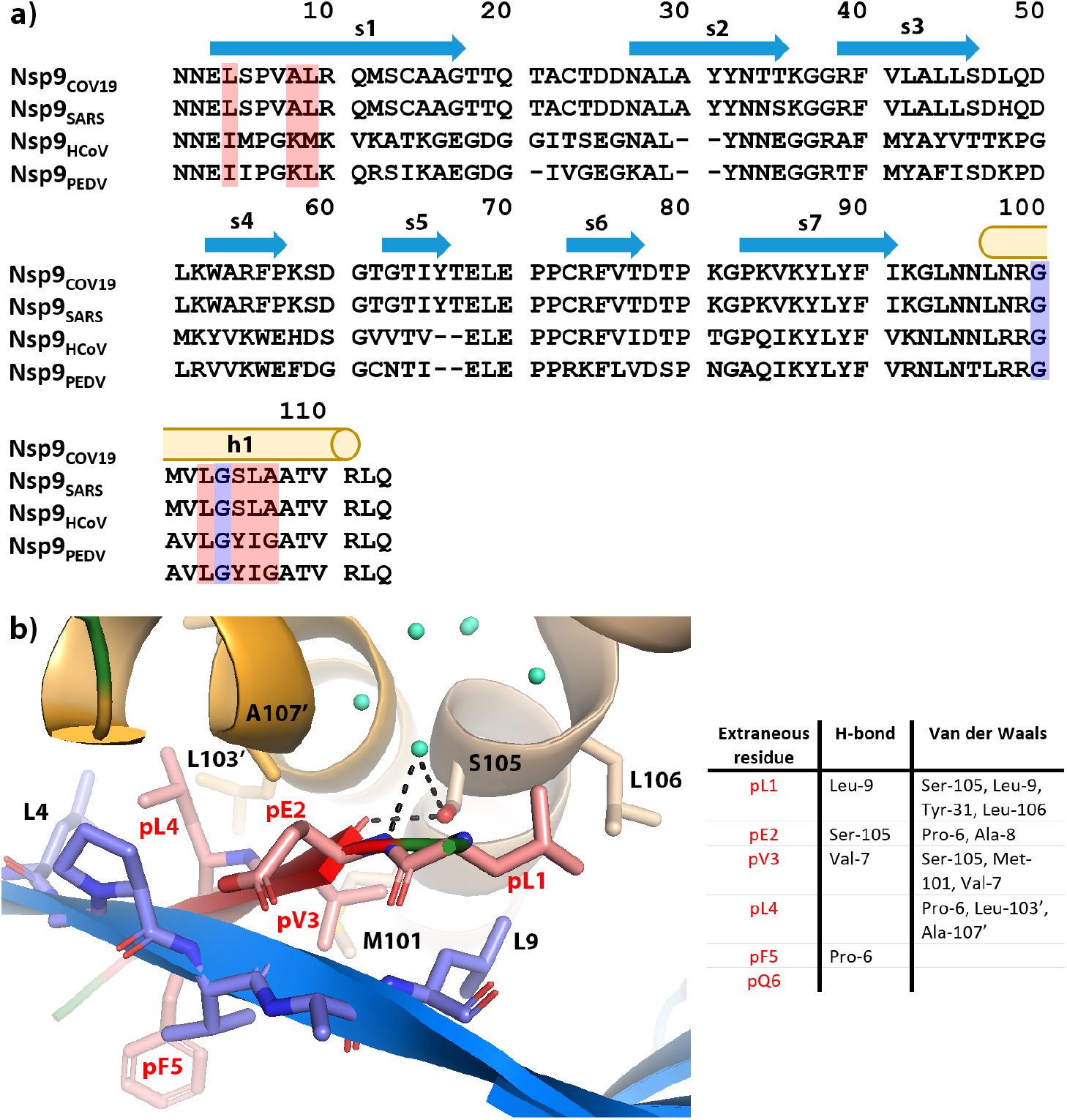
Sequence conservation within Nsp9 homologues. Sequence alignment for viral Nsp9 proteins encoded by SARS-CoV-2, SARS-CoV, Human coronavirus 229E and Porcine epidemic diarrhea virus. The extent of secondary structural elements observed in the 3C-Nsp9_COV19_ structure are shown and labelled above. The GxxxG motif residues are highlighted with *purple* and those making up the extraneous peptide binding site in *pink*. **b)** Cartoon and stick representation of the peptide-binding site observed in 3C-Nsp9_COV19_. The sidechains of the extraneous 3C residues on one side of the paired helical-interface are displayed with carbon atoms colored *pink*. Nearby residues that make up the binding site are displayed and labelled and listed in the accompanying contacts table.

Most residues involved in protein-binding within the hydrophobic cavity and the structural changes needed to accommodate them appear broadly conserved amongst other Nsp9 viral homologues (*red* highlights in Fig. 5A). The main exception to this is Ser-105, which is a Tyrosine in the distantly related Nsp9_HCoV_ and Nsp9_PEDV_ proteins (Ponnusamy et al., 2008), (Zeng *et al.*, 2018). However, the N-terminal interface β-strand in these homologues is known to be involved in interface re-organisation of the subunits (Ponnusamy et al., 2008) and thus denotes other structural differences at this site.

## Discussion

Here we describe the structure of the recombinantly expressed Nsp9_COV19_ as part of a global effort to characterise the virus causing a current global pandemic. Nsp9 is important for virulence in SARS-CoV (Miknis et al., 2009). It remains to be understood whether Nsp9_COV19_ plays a similar role in SARS-CoV-2, however the 97% sequence identity suggests a high degree of functional conservation. The CoV Nsp9 proteins are seemingly obligate dimers comprising a unique fold that associates via an unusual α-helical GxxxG interaction motif. The integrity of this motif is considered important for viral replication (Miknis et al., 2009), leading to a proposal that disruption of the unusual dimer interface impacts on RNA binding and function (Hu et al., 2017). Mutation of the same interaction motif in the porcine delta coronavirus Nsp9_PDCoV_ also disrupted nucleotide binding capacity (Zeng *et al.*, 2018).

We describe the ability to produce homogenous Nsp9_COV19_ which purifies as an obligate dimer, consistent with other Nsp9 proteins. Our preliminary nucleotide binding assays brought into question the RNA binding capacity of Nsp9_COV19_. The structure of the Nsp9_COV19_ showed conservation of the unique Nsp9 fold when compared with homologues from SARS (Egloff *et al.*, 2004) (Sutton *et al.*, 2004). Indeed, the topological fold was conserved as was the Nsp9 specific α-helical GxxxG dimerisation interface. This α-helical interface is encircled by hydrophobic residues but the interface includes considerable cavities as observed previously (Egloff *et al.*, 2004). We made a serendipitous discovery in our 3C-Nsp9_COV19_ structure, whereby the hydrophobic cavity captured the 3C cleavage sequence LEVL. The extraneous residues were tightly bound on all sides within the site situating themselves proximal to the conserved GxxxG motif. Coordination of the 3C sequence induced changes within interfacing residues, serving to both restructure key structural elements and cause a modest shift in subunit orientation.

At this stage it is unclear whether the bound residues within our structure have any bearing on the physiological function of Nsp9_COV19_. In the first instance this would seem unlikely, however our sequence is that of a rhinoviral 3C protease site and the SARS-CoV main protease cleaves consensus sequences following an LQ sequence (Zhu *et al.*, 2011). The bound 3C-LE residues have hallmarks of the LQ motif and are proximal to the highly conserved GxxxG motif. There are no obvious structural features to preclude, or select for, a Glu to Gln substitution within our bound sequence. Notably the M^pro^ cleavage sequence occurs at multiple points throughout the CoV genome as the majority of viral proteins are released by its activity, thus it remains possible that Nsp9 may associate with unprocessed viral polyproteins retaining them near the viral RNA. Within our structure Met-101 provides contacts with the bound valine sidechain but the presence of Asn-33 nearby may also accommodate residues such as lysine at this position. Peptide-binding assays will need to be developed to rigorously assess such possibilities.

In summary we have established a protocol for the production and purification of SARS-CoV-2 Nsp9 protein. We determined the structure of the Nsp9_COV19_ and described the conservation of the unique fold and dimerisation interface identified previously for members of this protein family. We also determined structure of Nsp9_COV19_ in complex with a 3C sequence, although the significance of this is yet to be established. The structures we describe here could potentially be utilised in drug screening and targeting experiments to disrupt a dimer interface known to be important for coronavirus replication.

## Materials and Methods

### Protein production, crystallisation, structure determination, and refinement

Synthetic cDNA for the Nsp9_COV19_ protein was cloned into pET28-LIC expression vector bearing an N-terminal His-tag with a Rhinovirus 3C protease cleavage site (MAHHHHHHSAA****LEVL****FQGPG). The plasmid was transformed into *E. coli* BL21(DE3) cells which were grown in Luria Broth at 37°C until reaching an Absorbance at 600nm of ~1.0 before being induced with 0.5mM Isopropyl β-d-1-thiogalactopyranoside for 4 hours. Cells were harvested in 20mM HEPES pH 7.0, 150mM NaCl, 20mM Imidazole, 2mM MgCl_2_ and 0.5mM TCEP and frozen until required. Lysis was achieved by sonicating the cells in the presence of 1mg of Lysozyme and 1mg of DNAase on ice. The lysate was then cleared by centrifugation at 10,000xg for 20 minutes and loaded onto a nickel affinity column. Bound protein was washed extensively with 20 column volumes of 20mM HEPES pH 7.0, 150mM NaCl, 0.5mM TCEP before being eluted in the same buffer with the addition of 400mM Imidazole. For His-tag removal samples were incubated with precision 3c protease overnight at 4°C. All samples were subjected to gel filtration (S75 16/60; GE Healthcare) in 20mM HEPES pH 7.0, 150mM NaCl before being concentrated to 50mg/mL for crystallization trials.

Nsp9_COV19_ crystallized in 2.0-2.2M NH_4_SO_4_ and 0.1M Phosphate-citrate buffer pH 4.0. Crystals of the His-tag samples grew with rectangular morphology in space group P4_1_22, however if the His-tag was removed the crystals grew in space group P6_1_22 form with hexagonal morphology. All diffraction data were collected at the Australian synchrotron’s MX2 beamline at the Australian synchrotron (Aragao *et al.*, 2018) (see Table 1 for details). Data were integrated in XDS (Kabsch, 2010), processed with SCALA, phases were obtained through molecular replacement using PDB 1QZ8 (Egloff *et al.*, 2004) as a search model. Subsequent rounds of manual building and refinement were performed in Coot (Casanal *et al.*, 2019) and PHENIX (Liebschner *et al.*, 2019).

### Nucleotide binding assay

To examine the RNA-binding affinity, an 18-point serial dilution (212-0 μM) of Nsp9_COV19_ was incubated with 1 nM 5’-Fluorescein labelled 17mer poly-U single-stranded RNA (Dharmacon GE, USA) or 10mer PolyT single-stranded DNA (IDT, USA) in assay buffer (20mM HEPES pH 7.0, 150mM NaCl, 2mM MgCl_2_) at room temperature.

The assay was performed in 96-well non-binding black plates (Greiner Bio-One), with fluorescence anisotropy measured in triplicate using the PHERAstar FS (BMG) with FP 488-520-520 nm filters. The data was corrected using the anisotropy of RNA sample alone, then fitted by a one-site binding model using the Equation, A = (Amax [L])/(KD+[L]), where A is the corrected fluorescence anisotropy; Amax is maximum binding fluorescence anisotropy signal, [L] is the Nsp9_COV19_ concentration, and *K*_*D*_ is the dissociation equilibrium constant. Amax and *K*_*D*_ were used as fitting parameters and nonlinear regression was performed using GraphPad Prism. Measurements were taken after 60 minutes incubation between protein and RNA.

## Supplementary note

During manuscript preparation, the Center for Structural Genomics of Infectious Diseases (CSGID) deposited the structure of Nsp9_COV19_ (6W4B) to a resolution of 2.95Å

## End Matter

### Author Contributions and Notes

D.R.L, B.S.G, and J.R. designed the project and wrote the manuscript. D.R.L. cloned, purified and crystallised Nsp9_Cov19_ and refined the structures. R.C.N. performed the RNA binding assays. Diffraction data and models have been deposited at the protein databank (www.rcsb.org) with accession codes 6W9Q and 6WC1.

The authors declare no conflict of interest.

This article contains supporting information online.

## Acknowledgments

Funding for the work originated from the Australian Research Council Centre of Excellence for Advanced Molecular Imaging. This research was undertaken in part using the MX2 beamline at the Australian Synchrotron, part of ANSTO, and made use of the Australian Cancer Research Foundation (ACRF) detector. We thank A. Riboldi-Tunnicliffe for assistance with data collection and J. Whisstock and G. Watson for advice on the manuscript.

